# Involuntary orienting and conflict resolution during auditory attention: The role of ventral and dorsal streams

**DOI:** 10.1101/2020.01.24.917310

**Authors:** Hannah J Stewart, Dawei Shen, Nasim Sham, Claude Alain

## Abstract

Selective attention to sound object features such as pitch and location is associated with enhanced brain activity in ventral and dorsal streams, respectively. We examined the role of these pathways in involuntary orienting and conflict resolution using functional magnetic resonance imaging (fMRI). Participants were presented with two tones that may share, or not, the same non-spatial (frequency) or spatial (location) auditory features. In separate blocks of trials, participants were asked to attend to sound frequency or sound location and ignore the change in the task-irrelevant feature. In both attend-frequency and attend-location tasks, response times were slower when the task-irrelevant feature changed than when it stayed the same (involuntary orienting). This behavioural cost coincided with enhanced activity in the prefrontal cortex and superior temporal gyrus (STG). Conflict resolution was examined by comparing situations where the change in stimulus features was congruent (both features changed) and incongruent (only one feature changed). Participants were slower and less accurate for incongruent than congruent sound features. This congruency effect was associated with enhanced activity in the prefrontal cortex, and was greater in the right STG and medial frontal cortex during the attend-location than during the attend-frequency task. Together, these findings do not support a strict division of ‘labour’ into ventral and dorsal streams, but rather suggest interactions between these pathways in situations involving changes in task-irrelevant sound feature and conflict resolution. These findings also validate the Test of Attention in Listening task by revealing distinct neural correlates for involuntary orienting and conflict resolution.

## Introduction

To navigate successfully in complex auditory environments, the listener needs to identify ***what*** different sounds are and ***where*** they are coming from (Alain & Arnott, 2000; Alain, Arnott, Hevenor, Graham, & Grady, 2001). Evidence from animal studies (Lomber & Malhotra, 2008; Rauschecker, 1998; Rauschecker & Tian, 2000; Romanski et al., 1999), as well as from functional magnetic resonance imaging (fMRI) studies (Alain et al., 2001; Alain, He, & Grady, 2008; Degerman, Rinne, Salmi, Salonen, & Alho, 2006; Leung & Alain, 2011; Maeder et al., 2001) suggest that ventral and dorsal brain areas play an important role in identifying and locating sounds in the environment, respectively. Human lesion studies have shown double dissociations between non-spatial and spatial auditory processing in patients with left and right hemisphere damage (Clarke, Bellmann, Meuli, Assal, & Steck, 2000; Clarke et al., 2002; Clarke & Thiran, 2004; Zundorf, Lewald, & Karnath, 2016). Meta-analyses of auditory fMRI studies provide further support for ventral and dorsal pathways involved in processing the sound object identity and location, respectively (Alho, Rinne, Herron, & Woods, 2014; Arnott, Binns, Grady, & Alain, 2004).

While a number of imaging studies have investigated the dual pathway theory with varying types of auditory stimuli and tasks (e.g., noise bands with center frequencies, Alain et al., 2001; dichotic vowel identification, Du et al., 2015; and 3-talker sentences, Hill & Miller, 2010), there have been few investigations into the auditory ventral and dorsal pathways with regards to specific attentional constructs (e.g., Mayer et al., 2006; Orr & Weissman, 2009). The few that do explore different attentional constructs assess each construct in isolation despite multiple attentional constructs being used in parallel in real life. Zhang, Barry, Moore & Amitay (2012) have developed a test, the Test of Attention in Listening (TAiL), based on two of Posner and Petersen’s (1990; Petersen & Posner, 2012) Attention Network constructs. During auditory attention tasks TAiL can isolate involuntary orienting (attending to a task-irrelevant property/object) and executive control (completing the dominant task while also performing a subdominant task, e.g., conflict resolution).

Orienting to auditory information can occur voluntarily via cues or be triggered involuntarily by unexpected novel sound events. Cues, whether valid or invalid, exert strong effects on behaviour and cortical responses (Mayer et al., 2009; Posner & Petersen, 1990). The TAiL paradigm does not involve cues and so focuses on involuntary orienting. This attention construct has been widely investigated through auditory oddball paradigms where an infrequent uncued stimuli change occurs (e.g., loudness, pitch, location) to elicit attentional capture. However, in the TAiL paradigm the frequency (pitch) and location changes are subtle and occur every trial. Neuroimaging of such a paradigm is novel.

Two papers using auditory stimuli similar to ourselves, but in an oddball task, compared cortical areas activated when involuntarily orienting to task-irrelevant stimuli (loudness deviant) during a non-spatial (pitch – Alho et al., 2015) and spatial (location – Salmi et al., 2009) task-relevant paradigm. They found that the two paradigms showed overlapping areas of activation in the posterior superior temporal sulcus, middle temporal gyrus and middle frontal gyrus. However, there was stronger bilateral activation in large areas of the superior temporal gyrus/sulcus in the non-spatial task compared to the spatial task and in the ventromedial prefrontal area for the opposite contrast. Loudness was used to trigger involuntary orienting in both of the paradigms as evaluating the use of the dual pathways was not a goal of these sister papers. Therefore, it is still unclear whether involuntarily orienting to non-spatial and spatial auditory information will cause segregation or overlapping of the ventral and dorsal streams.

A classic methodology for testing conflict resolution involves contrasting incongruent with congruent trials as in the Stroop task (Stroop, 1935). Roberts and Hall (2008) conducted a meta-analysis of the visual Stroop task and found evidence of a common frontoparietal network, including the anterior cingulate cortex (ACC), inferior frontal gyrus (IFG), parietal lobe and anterior insula. They followed this meta-analysis up with the same group of participants completing the traditional visual Stroop task and an auditory version where listeners indicated whether the speaker’s voice was high or low in pitch, while ignoring semantic content (the words ‘high’, ‘low’ and ‘day’). The findings from the fMRI study were consistent with the results of their meta-analysis with a common frontoparietal network for both paradigms. However, they also observed extensive activity in the left lateral prefrontal cortex during the auditory Stroop task along with activity in the superior temporal sulcus; and the visual Stroop task showed greater activity in the right inferior frontal gyrus. These results suggest a supramodal conflict resolution network with further areas activated for the separate modalities. However, the pathways activated by auditory spatial and non-spatial conflict resolution tasks has not been investigated. Siemann, Herrmann & Galashan (2018) investigated this question with visual stimuli and found comparable networks were activated for visual spatial and non-spatial conflict resolution (dorsal fronto-parietal networks in the left middle frontal gyrus, superior parietal lobule and frontal eye fields). With additional ventral areas recruited specifically for non-spatial conflict resolution (left middle temporal gyrus, cuneus and lingual gyrus).

Behavioural studies using TAiL provide support that the task taps into these two attentional constructs of involuntary orientation and conflict resolution using auditory stimuli (Zhang et al., 2012; Stewart & Amitay, 2015). Using scalp-recordings of event-related potentials (ERPs), Stewart and colleagues (2017) showed distinct ERP modulations associated with involuntary orienting and conflict resolution. The comparison of distributed source analyses for involuntary orienting suggests a more dorsal source when the task-irrelevant sound feature was location than when it was frequency. Although this finding appears consistent with the dual pathway model of auditory attention, further research is needed to more precisely identify the brain areas associated with involuntary orienting and conflict resolution during the TAiL task.

This current study combines fMRI, with its advantageous spatial resolution, and the TAiL task to investigate the brain areas associated with involuntary orienting and conflict resolution. The paradigm used provides a novel approach to this question as it allows both attentional constructs to be investigated using the same auditory trials and without the sustained attention effects of an oddball paradigm. It is expected that optimum task performance may differentially involve ventral and dorsal streams to process non-spatial and spatial auditory stimulus features, respectively. As found by Alho et al. (2015), we anticipate that involuntary orienting will engage a similar attention network for both spatial and non-spatial tasks including the superior temporal sulcus/gyrus, middle temporal gyrus and middle frontal gyrus. Due to the paradigm differences between TAiL’s continuous subtle task-irrelevant changes and previous studies’ deviant salient task-irrelevant changes, it is unclear which cortical areas will show additional activity for spatial and non-spatial stimuli. Following meta-analyses of the dual pathway theory in auditory tasks (Arnott et al., 2004; Alho et al., 2014), additional inferior parietal lobe activity is predicted for spatial stimuli and middle superior temporal gyrus activity for non-spatial stimuli. Conflict resolution contrasts are predicted to show frontoparietal activation including the ACC, IFG and parietal lobe, as found by Roberts and Hall (2008) and Siemann et al. (2018). However, it is unclear whether larger supramodal or auditory-specific networks will be engaged for the spatial (e.g., frontal cortex) and non-spatial auditory stimuli (e.g., amodal: middle temporal gyrus; supramodal: cuneus and lingual gyrus).

## Method

### Participants

Seventeen right-handed participants aged 19 to 30 years (Mean = 24.74 years, SD = 3.43 years; 13 females and 4 males) were recruited through the Rotman Research Institute participant database. Inclusion criteria was normal hearing (thresholds below 25 decibel (dB) hearing level (HL) bilaterally at frequencies between 250 and 8000 Hz, inclusive, and a normal score (0-3) on the Quick Speech-in-Noise test (QuickSIN) (Killion, Niquette, Gudmundsen, Revit, & Banerjee, 2004). Exclusion criteria was any self-reported history of brain damage, brain surgery, history of language-related or attention-related conditions, autism spectrum disorders or any auditory system disorders. All methods were approved by the Research Ethics Board at Baycrest Health Sciences, and performed in accordance with the relevant guidelines and regulations of Toronto Academic Health Services Network. All participants signed informed consent prior to the experiment and received monetary compensation for their participation.

### Stimuli and Task

All tones were made up of sinusoids with a duration of 100 ms, gated on/off by 10 ms cos ramps, and were presented monaurally at about 85 dB sound pressure level (SPL) (root mean square) by means of circumaural, fMRI-compatible headphones (Avotec, Jensen Beach, FL), acoustically padded to suppress scanner noise by about 25 dB SPL. Tone frequency was randomly selected for each participant from the range 476.18 - 6187.50 Hz with at least 2.1 equivalent rectangular bandwidths (ERBs; ∼ 4 semitones) between the trial’s tones, therefore making it well within the listener’s ability to discriminate between different frequencies (Jensen & Neff, 1993). The inter-trial-interval varied randomly between 1000 and 4000 ms (1000 ms steps, rectangular distribution), whilst the inter-stimulus interval (ISI) was set at 300 ms (Figure 1).

**Figure 1.**
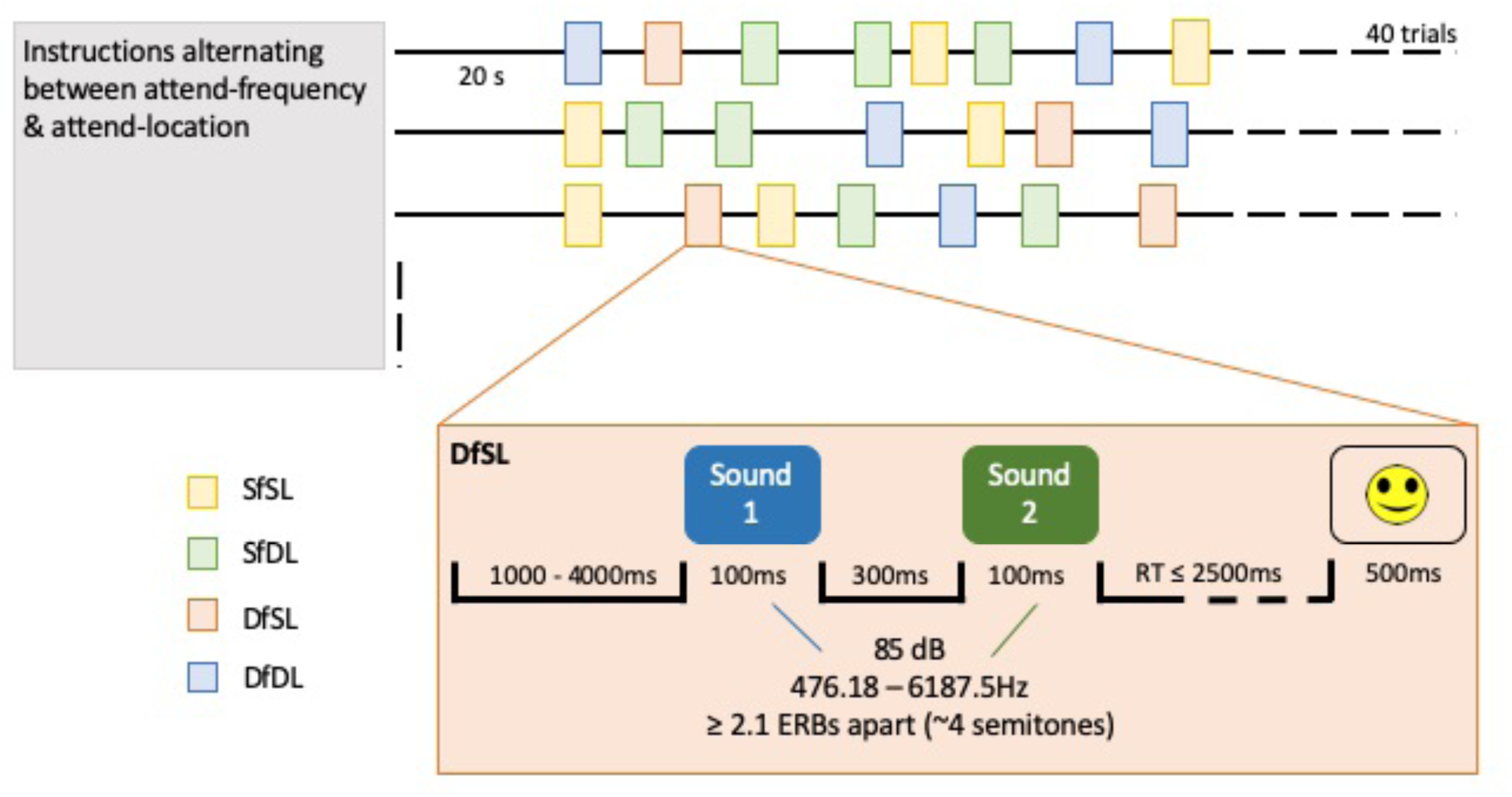
Paradigm schematics of runs and a single (DfSL) trial. Twelve runs were made up of alternating attend-frequency and attend-location TAiL tasks, counterbalanced across participants. Within each run 40 trials were presented consisting of 10 each of SfSL, SfDL, DfSL, DfDL trials randomized for each participant.

In both attend-frequency and attend-location tasks, the stimuli and paradigm remained the same, and only the instructions to the participants changed. In each trial, participants heard a tone pair where the individual tones were either the same or different in frequency and/or spatial location (ear presentation) (SfSL – same frequency and same location; DfSL – different frequency same location; SfDL – same frequency different location; DfDL – different frequency different location.). In both attend-frequency and attend-location tasks, the listener had to indicate via a button press if the task-relevant sound feature (i.e., the location of the two tones in the attend-location task) were same or different, while ignoring the task-irrelevant sound feature (i.e., the frequency of the two tones in the attend-location task). A full description of the task and stimuli can be found in Zhang et al (2012).

Listeners were asked to respond as fast and as accurately as possible after the second tone. Responses less than 200 ms and more than 2500 ms were excluded from further analysis in case of premature responding and interruption of performance. Participants’ responses were registered using an fMRI-compatible response pad (Lightwave Technologies, Surrey, SC, Canada). The left index finger was used for a ‘same’ response and the right index finger for a ‘different’ response. As soon as the listener responded to each trial (attend-frequency task group mean: 610.40 ms, SD: 268.86 ms, range: 208.50-1979.00 ms); attend-location task group mean: 598.80 ms, SD: 263.31 ms, range: 221.40-1985.70 ms) visual feedback was provided for 500 ms. If they answered correctly, a smiley face was displayed. If they answered incorrectly, the same face was shown with a sad expression. TAiL stimuli were automated and presented using Presentation software (version 16).

From the attend-frequency and attend-location tasks, three measures were calculated: *baseline, involuntary orienting* (the behavioural cost associated with a change in the task-irrelevant sound feature), and *conflict resolution* (the behavioural cost associated with processing conflicting sound feature). The *baseline* measure is calculated for each task type from trials where both sound features, frequency and location, remain constant (i.e., SfSL trials). The *involuntary orienting* measure is calculated as the difference in RT and accuracy between conditions when the task-irrelevant feature was different between Tone 1 and Tone 2 and when it was constant (see Table 1). The *conflict resolution* measure is calculated as the difference between trials where both auditory features remain constant or change together (congruent trials, SfSL and DfDL), compared with trials where only one feature changes (incongruent trials, DfSL and SfDL).

**Table 1:**
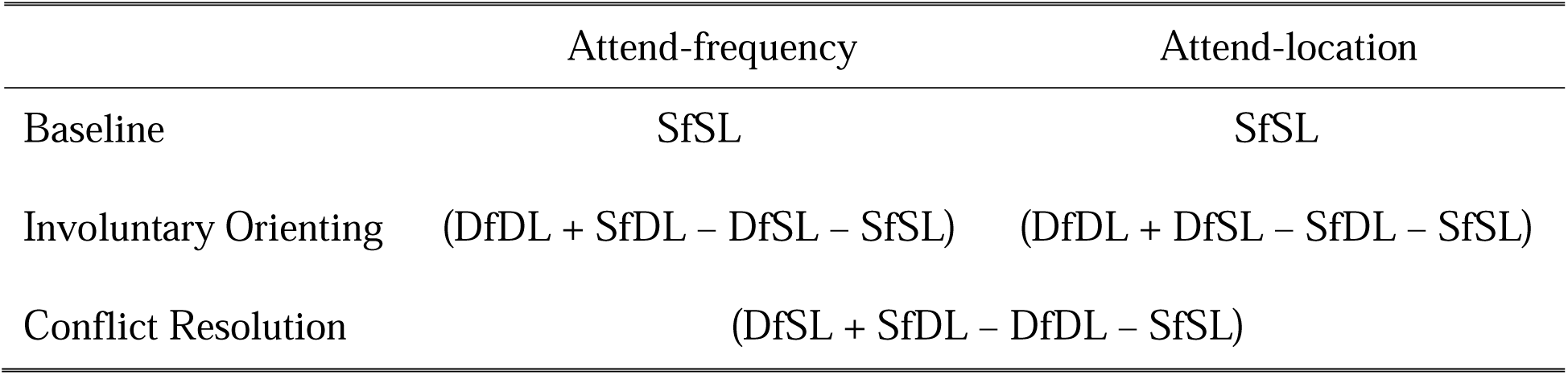
Calculations for the TAiL outcome measures

### Procedure

Before entering the scanner, the participants completed a practice block for each of the attend-frequency and attend-location tasks. Each practice block involved five trials, accompanied by on-screen instructions. Participants had to reach 60% accuracy or more to move onto the full testing blocks in the scanner. Each run consisted of 40 trials - 10 each of SfSL, DfSL, SfDL, DfDL - presented in a random order for each participant. Both tasks (attend-frequency and attend-location) had six runs providing a total of 40 trials per task per participant (Figure 1). The order of the task types was counterbalanced across participants and alternated across the runs, allowing regular rests for the participants. At the start of each run, participants were instructed of the task type and to respond as fast and as accurately as possible, and visual feedback regarding the listener’s performance was provided after each trial throughout. The total testing/recording time lasted around 45 minutes.

### Behavioural analysis

Reaction times (RTs) from correct trials, and accuracy (% correct) were used in the analysis. Repeated-measures ANOVAs with the task-relevant and task-irrelevant features as within-subjects factors were run for each TAiL task.

#### fMRI scanning and data analysis

Participants were scanned using a research-dedicated whole-body 3.0 T MRI system (Siemens Tim Trio – 3T software level Syngo MR 2006 VB13T) with a standard 12-channel quadrature bird-cage head coil. Structural T1 weighted anatomical volume were obtained at the midpoint of the experiment using SPGR (axial orientation, TR = 2000 ms, TE = 2.63 ms, FOV = 256 mm, slice thickness = 1 mm) for co-registration with the functional images and to ensure that there were no significant brain abnormalities in any of the participants.

Each functional scan sequence began with a 20 second period where no stimuli were presented. We used an event-related design with a continuous analyses image acquisition. Functional imaging was performed to measure brain activation by means of the BOLD effect (Ogawa, Lee, Kay, & Tank, 1990). Functional data were acquired using a whole head T2*-weighted echo-planar image sequence (TE: 30 ms; TR: 2 s; flip angle: 70°; 30 oblique axial slices with interleaved acquisition, 3.125mm×3.125mm×5mm voxel resolution; FOV: 200 mm; AM: 64×64). Physiological respiratory and cardiac waveforms were recorded from the bellows and photoplethysmograph peripherals on the scanner, respectively, using LabView (National Instruments, Austin TX).

In each run, the first ten scans were discarded to allow the magnetization to reach steady state. fMRI data was pre-processed and analyzed using Analysis of Functional Neuroimages software (Version AFNI_2011_12_21_1014) (Cox, 1996, 2012). In the pre-processing stage, the RETROICOR technique (Glover, Li, & Ress, 2000) was used to perform physiological noise correction. By performing a slice timing correction, all slices can be aligned to the time of the acquisition for the center slice (For the Siemens 3T Trio Scanner at Baycrest, slice 1 is the center slice for each TR). To perform rigid body motion correction, for each run, images acquired at each point in the series were aligned volumetrically, using the 3dvolreg plugin for AFNI, to a reference image acquired during the scanning session. We chose the 51^st^ scan of the middle run (the 3^rd^ run in the present study) as a reference scan because the 51^st^ scan is usually very reliable for most datasets. Note that the 51^st^ scan corresponds to subbrick 50 because the 1^st^ scan is subbrick 0. The head motions are captured by 6 motion parameters: roll = rotation about the I-S axis; pitch = rotation about the R-L axis; yaw = rotation about the A-P axis; dS = displacement in the Superior direction; dL = displacement in the left direction; and dP = displacement in the Posterior direction. The maximum rotation is less than 1.5 degree, and the peak range of displacement was less than 1.5 mm for all participants. The co-registration results were also checked visually for additional quality control. To effectively compare fMRI data across participants, baseline normalization was performed by calculating the percent change. The percent signal change is calculated for each participant on a voxel-by-voxel basis. The program 3dAutomask was used to create a mask image to specify what part of the image is of the brain and what part is not. It can eliminate noise outside the brain and reduce the number of voxels.

The program 3dDeconvolve, which provides deconvolution analysis of fMRI time series data on the voxel by voxel basis, was performed on the pre-processed data by using a linear fitting. The shape of the response was modelled as a “gamma” function time-locked on trial onset for each stimulus condition. Only trials where participants responded correctly were included in the event-related analysis. For each participant we used all runs to create activation maps for each of the four conditions: SfSL DfSL SfDL DfDL. These activation maps were then spatially normalized to a Montreal Neurological Institute (MNI) template. We decreased spatial noise variance by convolving the fMRI brain volume to a 6 mm FWHM smoothness using 3dFWHM. The images were also de-trended by fitting a 3^rd^ order Legendre polynomial at each voxel, and regressing it out of the time series.

AFNI’s general linear model (GLM) (Winkler, Ridgway, Webster, Smith, & Nichols, 2014) was then used to perform second level group analysis and create maps to identify statistically significant (threshold *p* < 0.005, *F* ≥ 3.252) group effect clusters in the various contrasts. Monte Carlo simulations with AlphaSim in AFNI was run to perform Family-wise Error Correction for multiple comparisons (uncorrected p-value = .005; corrected p-value = .03; minimum cluster size = 12; cluster connection radius = 6.67 mm); only clusters larger than 586 uL were selected. These single-participant univariate maps were then warped to MNI space, using the EPI to MNI transformation that was previously computed for each participant’s data. The program of Surface Mapping with AFNI (SUMA) was used to display the images.

## Results

### Baseline

The group mean accuracy and reaction times (RTs) are presented in Table 2. In trials with no distracting or conflicting auditory information (i.e., SfSL – same frequency and same location) performance was comparable between the two tasks (attend-frequency and attend-location) for RT, *t*(16) = −1.35, *p* = .20, and accuracy, *t*(16) = 1.45, *p* = .17. This result suggests that differences, behavioural and cortical, in involuntary orienting and conflict resolution between the two tasks cannot be easily accounted for by differences in task difficulty.

**Table 2:** RT and accuracy for the four trial types in each TAiL task.

| Conditons: |  | RT (ms) |  | Accuracy (%) |  |
| --- | --- | --- | --- | --- | --- |
|  |  | Mean | SEM | Mean | SEM |
| Attend-frequency | SfSL | 492.41 | 23.51 | 98.60 | 0.79 |
|  | SfDL | 675.30 | 38.30 | 91.95 | 2.67 |
|  | DfSL | 627.38 | 37.62 | 94.80 | 1.60 |
|  | DfDL | 646.55 | 37.93 | 94.54 | 1.36 |
| Attend-location | SfSL | 517.06 | 25.44 | 97.75 | 0.84 |
|  | SfDL | 625.93 | 31.73 | 91.84 | 1.98 |
|  | DfSL | 657.89 | 27.26 | 89.59 | 1.91 |
|  | DfDL | 594.31 | 33.8 | 97.08 | 0.97 |

### Involuntary orienting

#### Behavioural results

A significant effect of distraction (i.e., when the task-irrelevant sound features were different vs. the same) was found in each TAiL task. In the attend-frequency task, participants were significantly slower at responding to trials where the task-irrelevant location of the sounds changed (SfDL – same frequency and different location; DfDL – different frequency and different location) compared to when they stayed constant (SfSL – same frequency and same location; DfSL – different frequency and same location) (RT: *F*(1, 16) = 66.64, *p* < .001, η_p_^2^ = .81) (Figure 2A). However, they showed no difference in accuracy (*F*(1, 16) = 36.27, *p* = .097, η_p_^2^ = .16) (Figure 3A). In the attend-location task, listeners were significantly slower and less accurate at responding to trials where the irrelevant frequency of the sounds changed (DfSL – different frequency and same location; DfDL – different frequency and different location) compared to when they stayed constant (SfSL – same frequency and same location; SfDL – same frequency and different location) (RT: *F*(1, 16) = 75.54, *p* < .001, η_p_^2^ = .83; accuracy: *F*(1, 16) = 8.12, *p* = .012, η_p_^2^ = .34) (Figures 2B and 3B).

**Figure 2.**
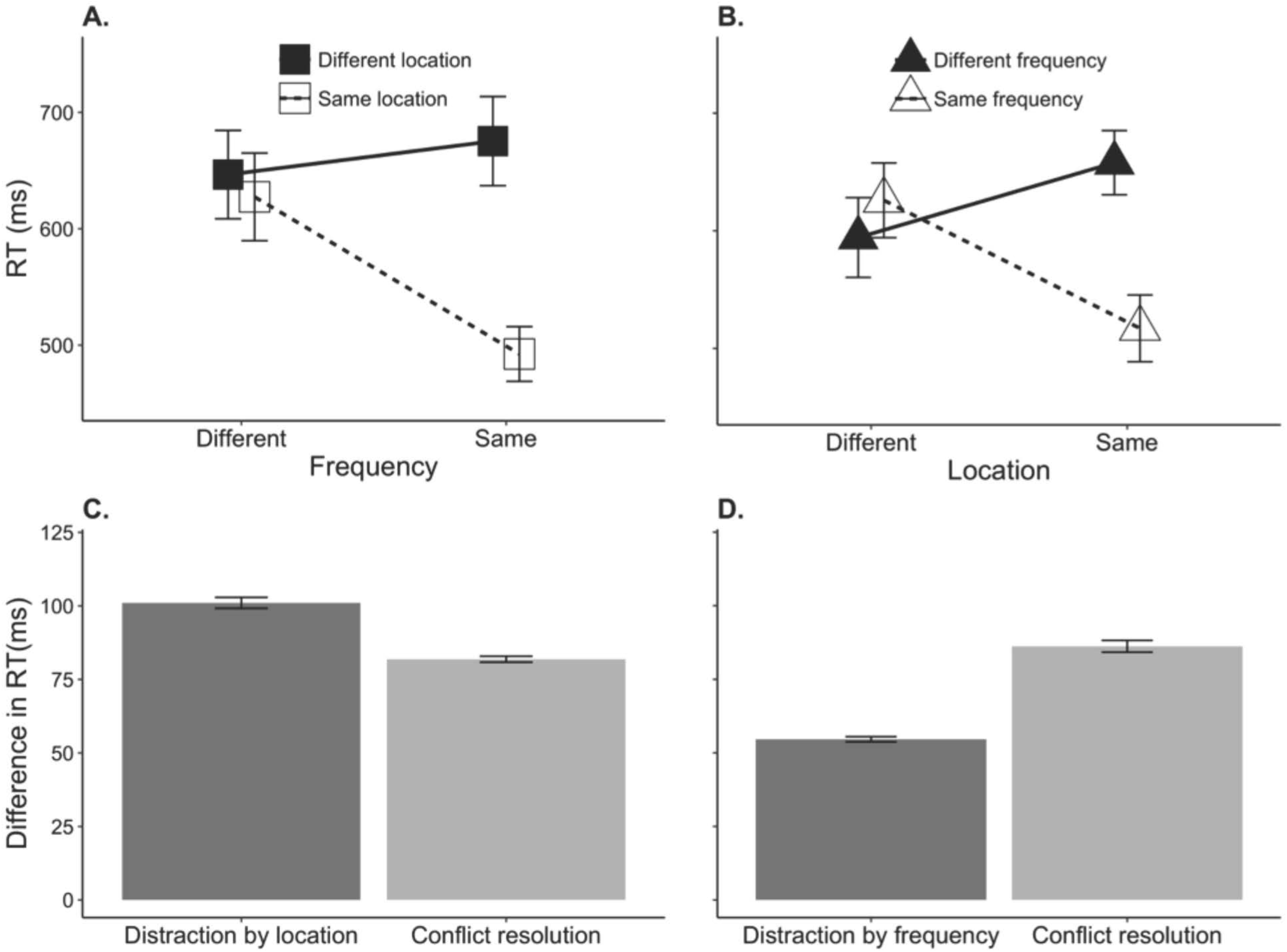
Group mean response times for the (A) attend-frequency and (B) attend-location tasks. Group mean measures of involuntary orienting, and conflict resolution for the (C) attend-frequency and (D) attend-location tasks. The error bars represent the standard error of the mean.

**Figure 3.**
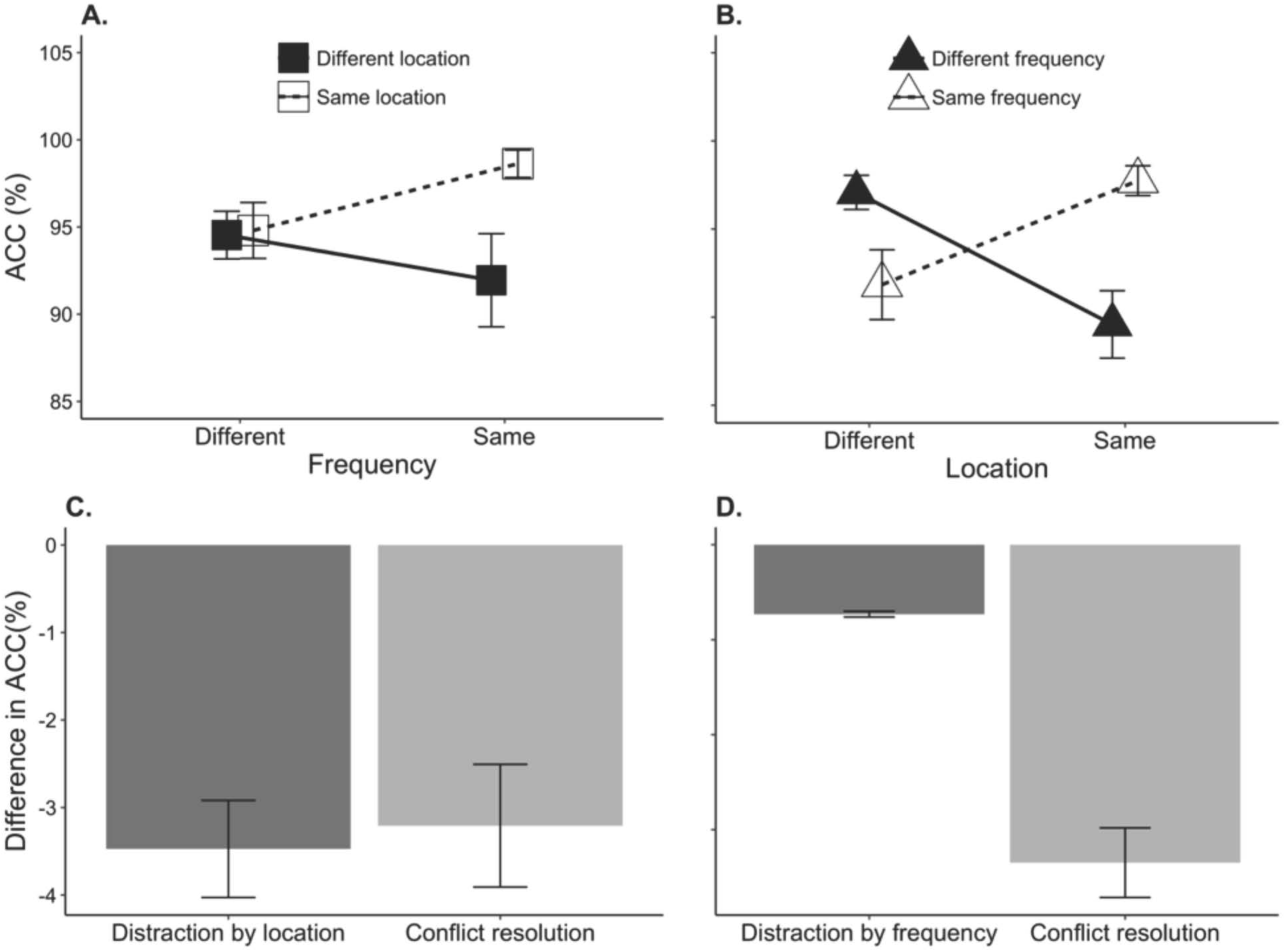
Group mean accuracy for the (A) attend-frequency and (B) attend-location tasks. Group mean measures of involuntary orienting, and conflict resolution for the (C) attend-frequency and (D) attend-location tasks. The error bars represent the standard error of the mean.

The involuntary orienting difference in RT was significantly lower in the attend-location task compared to the attend-frequency task, *t*(16) = 3.25, *p* = .005 (Figures 2C and D), but not for accuracy, *p* = .27 (Figures 3C and D).

To sum up, in both task, RTs were slower when the task-irrelevant feature changed than when it stayed the same, and this effect was greater when attention was focused on non-spatial (frequency) than spatial (location) auditory feature.

#### fMRI results

Figures 4 and 5 show activity associated with involuntary orienting during attend-frequency and attend-location, respectively. As for the behavioural data, the neural correlates were examined by contrasting BOLD responses when the task-irrelevant feature changed versus when the same task-irrelevant feature was repeated within the trial. When participants were instructed to focus attention on frequency, changes in task-irrelevant sound location were associated with increased BOLD signal bilaterally in superior frontal gyrus (SFG), the right inferior frontal gyrus (IFG), the left superior temporal gyrus (STG) and the left precentral gyrus (Table 3). Conversely, when participants were instructed to focus attention on sound location, changes in task-irrelevant sound frequency were associated with increased BOLD signal in right middle frontal gyrus (MFG), right IFG, bilateral STG, and bilateral IPL (Table 4).

**Table 3.** Attend-frequency task: Different location vs. same location (DfDL+SfDL-DfSL-SfSL) (*pthr* = .005, t = 3.252; corrected *p* < .05)

| Brain Regions | Peak MNI co-ordinate |  |  |  | t-values |
| --- | --- | --- | --- | --- | --- |
|  | BA | x | y | z |  |
| R inferior frontal gyrus (IFG) | 45 | 41 | 25 | -1 | 5.42 |
| L & R superior frontal gyrus (SFG) | 8 | 3 | 30 | 52 | 5.26 |
| L & R caudate and commissure |  | 1 | -26 | -2 | 3.46 |
| L inferior frontal gyrus (IFG) | 45 | -33 | 26 | -1 | 5.98 |
| L precentral gyrus | 6 | -28 | -6 | 68 | 3.68 |
| L superior temporal gyrus (STG) | 22 | -62 | -38 | 22 | 3.34 |

**Table 4.** Attend-location task: Different frequency vs. same frequency (DfDL+DfSL-SfDL-SfSL) (*pthr* = .005, t = 3.252; corrected *p* < .05)

| Brain Regions | Peak MNI co-ordinate |  |  |  | t-values |
| --- | --- | --- | --- | --- | --- |
|  | BA | x | y | z |  |
| R inferior frontal gyrus (IFG) | 9 | 50 | 12 | 29 | 3.26 |
| R middle frontal gyrus (MFG) | 9 | 52 | 35 | 25 | 3.75 |
| R superior temporal gyrus (STG) | 21 | 69 | -25 | -29 | 4.51 |
| R inferior parietal lobule (IPL) | 40 | 56 | -46 | 44 | 3.50 |
| L superior temporal gyrus (STG) | 42 | -64 | -25 | -4 | 3.48 |
| L inferior parietal lobule (IPL) | 40 | -49 | -42 | 57 | 4.06 |

**Figure 4.**
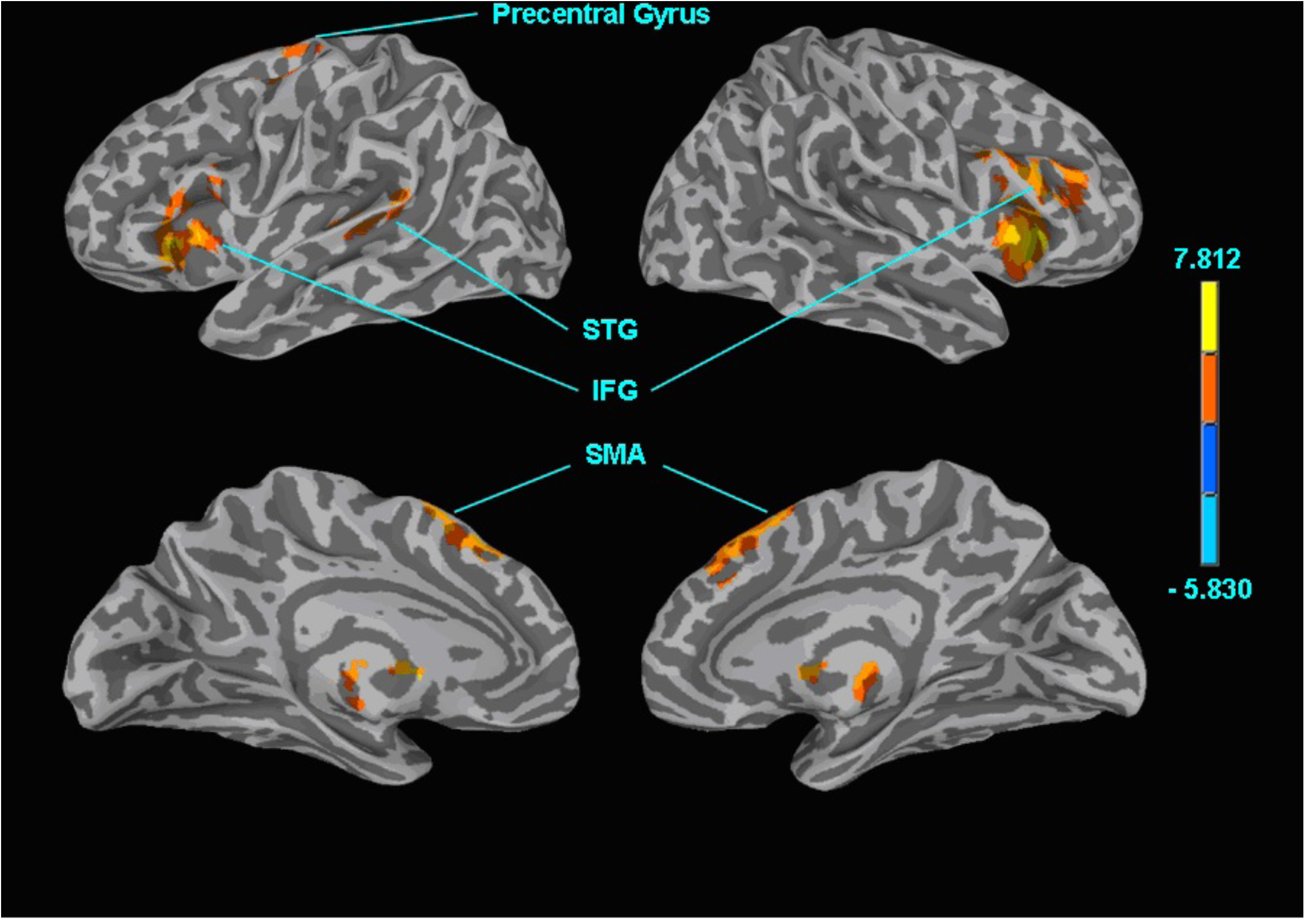
Attend-frequency task: Different Location vs. same location. Surface maps showing the effects of the irrelevant location feature while attending to the relevant frequency feature (i.e. involuntary orienting). Abbreviations: STG = superior temporal gyrus; SMA = supplementary motor area; IFG = inferior frontal gyrus.

**Figure 5.**
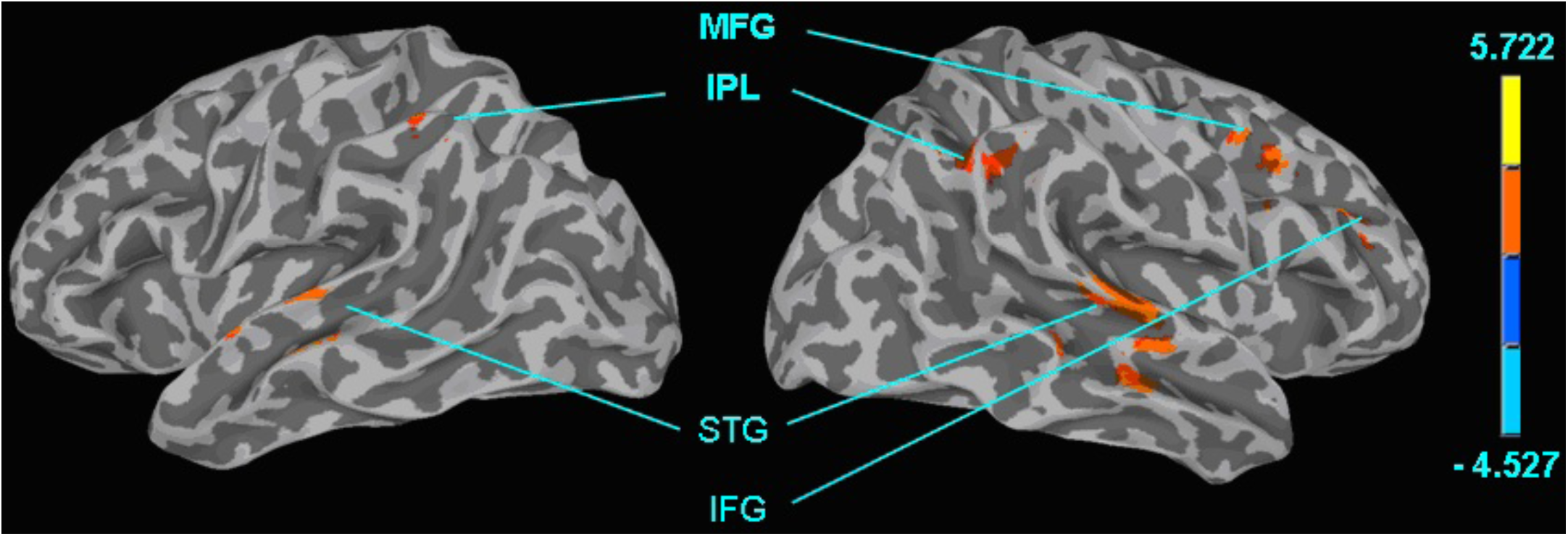
Attend-location task: Different frequency vs. same frequency. Surface maps showing the effects of the irrelevant frequency feature while attending to the relevant location feature (i.e. involuntary orienting). Abbreviations: MFG = middle frontal gyrus; IPL = inferior parietal lobule; STG = superior temporal gyrus; IFG = inferior frontal gyrus.

We also tested whether the changes related to involuntary orienting differed between the attend-frequency and the attend-location tasks. The attend-frequency task (SfDL-SfSL) minus attend-location task (DfSL-SfSL) revealed greater activity in the attend-location task in the right superior temporal gyrus (BA 22, Figure 6). That is, task-irrelevant changes in sound frequency was associated with enhanced activity in right STG. Although this difference in activation is relatively small, it suggests that the dorsal and ventral pathways are differentially recruited in situations designed to “trigger” involuntary orienting (Table 5).

**Table 5.**
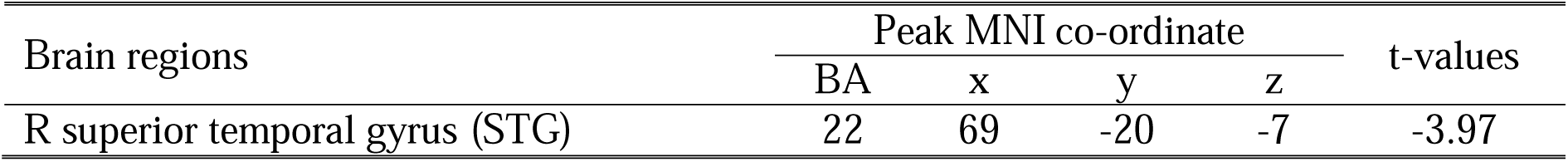
Attend-frequency vs. attend-location task: Involuntary orienting (SfDL+ SfSL – DfSL - SfSL) (*pthr* = .005, t = 3.252; corrected *p* < .05)

**Figure 6.**
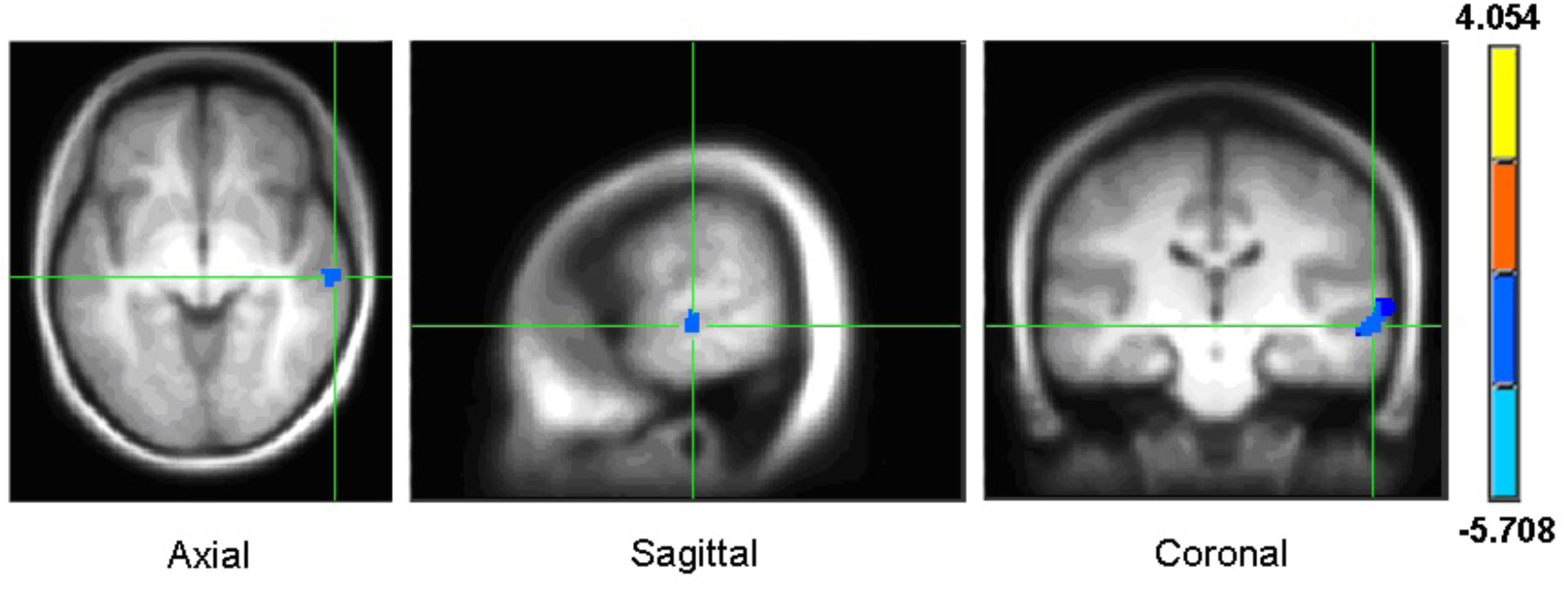
Contrast between the two involuntary orienting effects (attend-frequency > attend-location).

### Conflict resolution

#### Behavioural results

Evidence of *conflict resolution* was found in each TAiL task. In the attend-frequency task, participants were significantly slower and less accurate at responding to trials with incongruent sound features (SfDL – same frequency and different location; DfSL – different frequency and same location) compared to trials with congruent sound features (SfSL – same frequency and same location; DfDL – different frequency and different location) (RT: *F*(1, 16) = 49.95, *p* < .001, η_p_^2^ = .76; accuracy: *F*(1, 16) = 9.80, *p* = .006, η_p_^2^ = .38) (Figures 2A and 3A). Similarly, in the attend-location task, listeners were significantly slower and less accurate at responding to incongruent trials compared to congruent trials (RT: *F*(1, 16) = 113.54, *p* < .001, η_p_^2^ = .88; accuracy: *F*(1, 16) = 21.55, *p* < .001, η_p_^2^ = .57) (Figures 2B and 3B).

The *conflict resolution* difference in RT was comparable between the two tasks (attend-frequency and attend-location), *t*(16) = −0.11, *p* = .91 (Figures 2C and D), but the difference in accuracy was significantly smaller for the attend-frequency task, *t*(16) = 2.55, *p* = .022 (Figures 3C and D).

#### fMRI results

The neural activity associated with *conflict resolution* is shown in Figures 7 and 8. As for the behavioural data, the neural correlates were examined by contrasting BOLD responses when one task feature changed (incongruent trials) versus when both of the task features changed or stayed constant within the trial (congruent trials). When participants were instructed to focus attention on frequency, incongruent trials (DfSL+SfDL) generated a greater BOLD response than congruent (DfDL+SfSL) trials in several brain regions. These included bilateral SFG and IFG, MFG, and right anterior cingulate cortex, as well as right caudate and left paraphippocampal gyrus (Table 6).

**Table 6.** Attend-frequency task: Incongruent vs. congruent (DfSL+SfDL-DfDL-SfSL) (*pthr* = .005, t = 3.252; corrected *p* < .05)

| Brain Regions | Peak MNI co-ordinate |  |  |  | t-values |
| --- | --- | --- | --- | --- | --- |
|  | BA | x | y | z |  |
| R inferior frontal gyrus (IFG) | 47 | 52 | 22 | -5 | 3.73 |
| R & L superior frontal gyrus (SFG) | 6 | 3 | 17 | 53 | 3.73 |
| R cingulate gyrus | 23 | 20 | -5 | 30 | 3.83 |
| R & L culman | 35 | 18 | -27 | -20 | 4.04 |
| R caudate head |  | 16 | 29 | 1 | 3.60 |
| R caudate tail |  | 29 | -40 | 21 | 3.45 |
| L precentral gyrus | 9 | -45 | 9 | 34 | 4.15 |
| L inferior frontal gyrus (IFG) | 13 | -37 | 22 | -3 | 5.02 |
| L parahippocampal gyrus | 19 | -37 | -48 | 2 | 3.90 |

**Figure 7.**
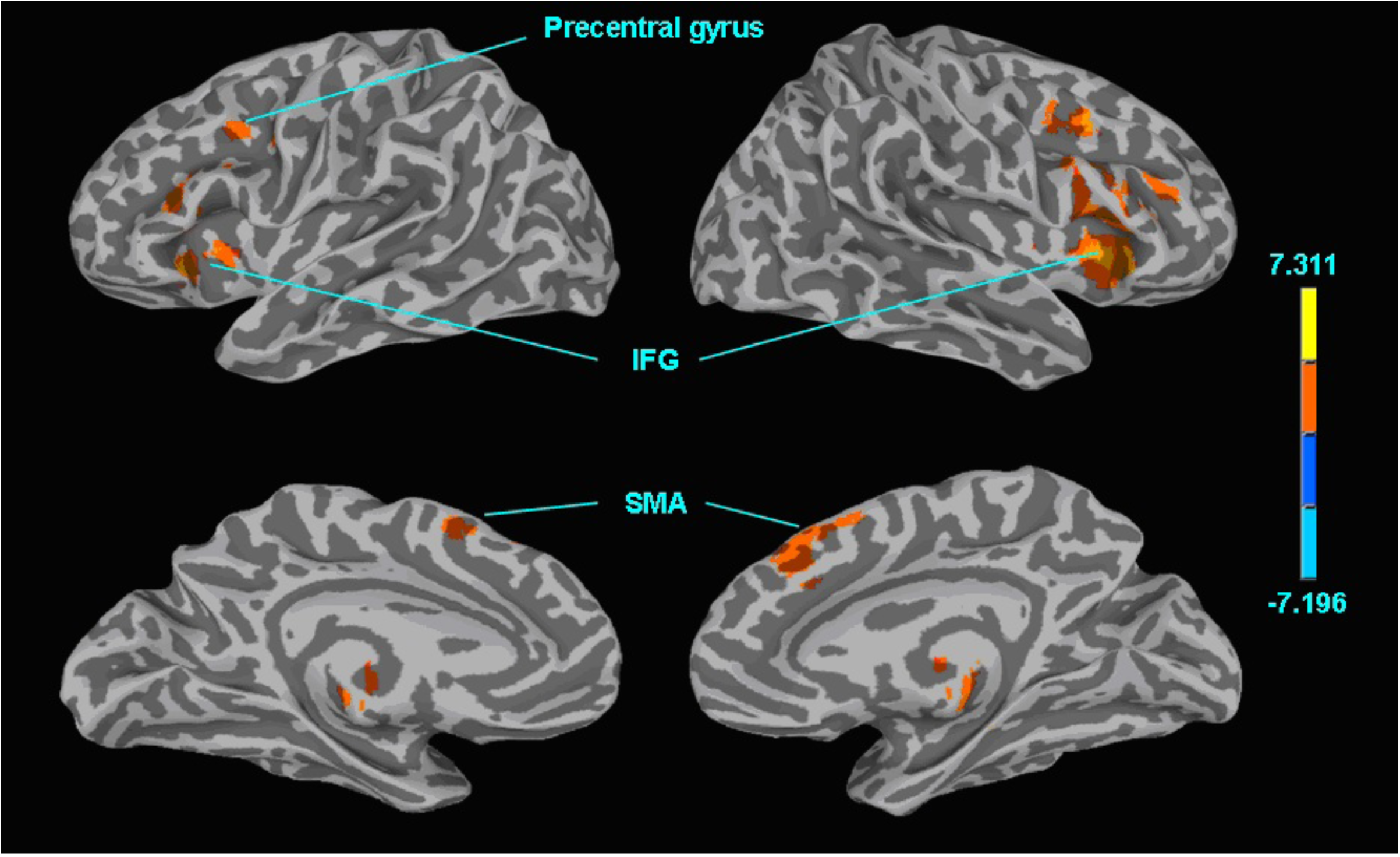
Attend-frequency task: Incongruent vs. congruent. Surface maps showing the effects of incongruency (DfSL+SfDL-DfDL-SfSL) while attending to the relevant frequency feature (i.e., conflict resolution). Abbreviations: IFG = inferior frontal gyrus; SFG = superior frontal gyrus.

**Figure 8.**
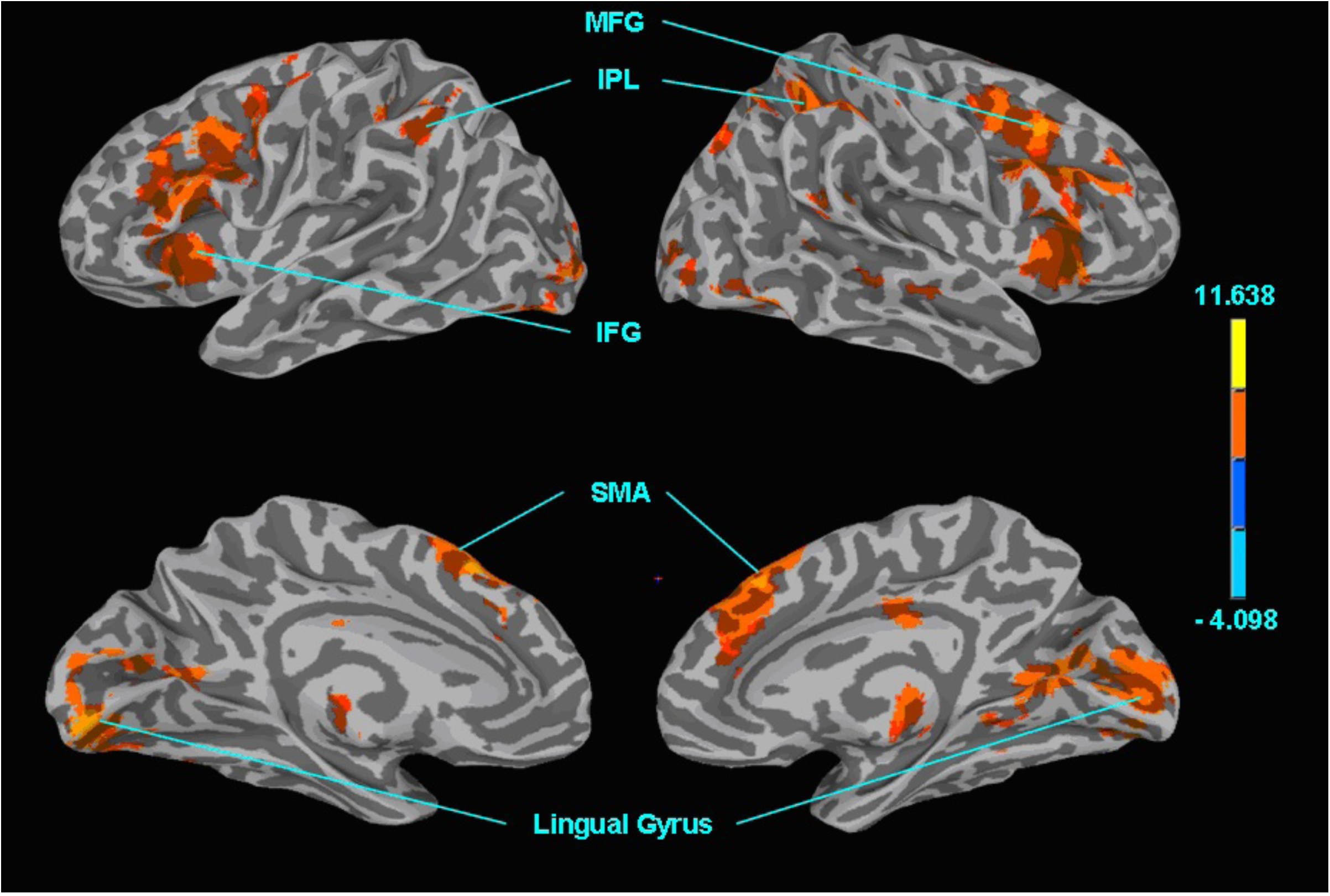
Attend-location task: Incongruent vs. congruent. Surface maps showing the effects of incongruency (DfSL+SfDL-DfDL-SfSL) while attending to the relevant location feature (i.e., conflict resolution). Abbreviations: MFG = middle frontal gyrus; IPL = inferior parietal lobule; IFG = inferior frontal gyrus; SMA = supplementary motor areas; STG = superior temporal gyrus.

Conversely, when participants were instructed to focus attention on sound location, the incongruent trials, relative to congruent trials, were associated with increased activation in bilateral SFG, right STG, left IFG and left IPL, cerebellum, as well as right MFG, right precuneus, right MTG, and bilateral cingulate (Table 7).

**Table 7.**
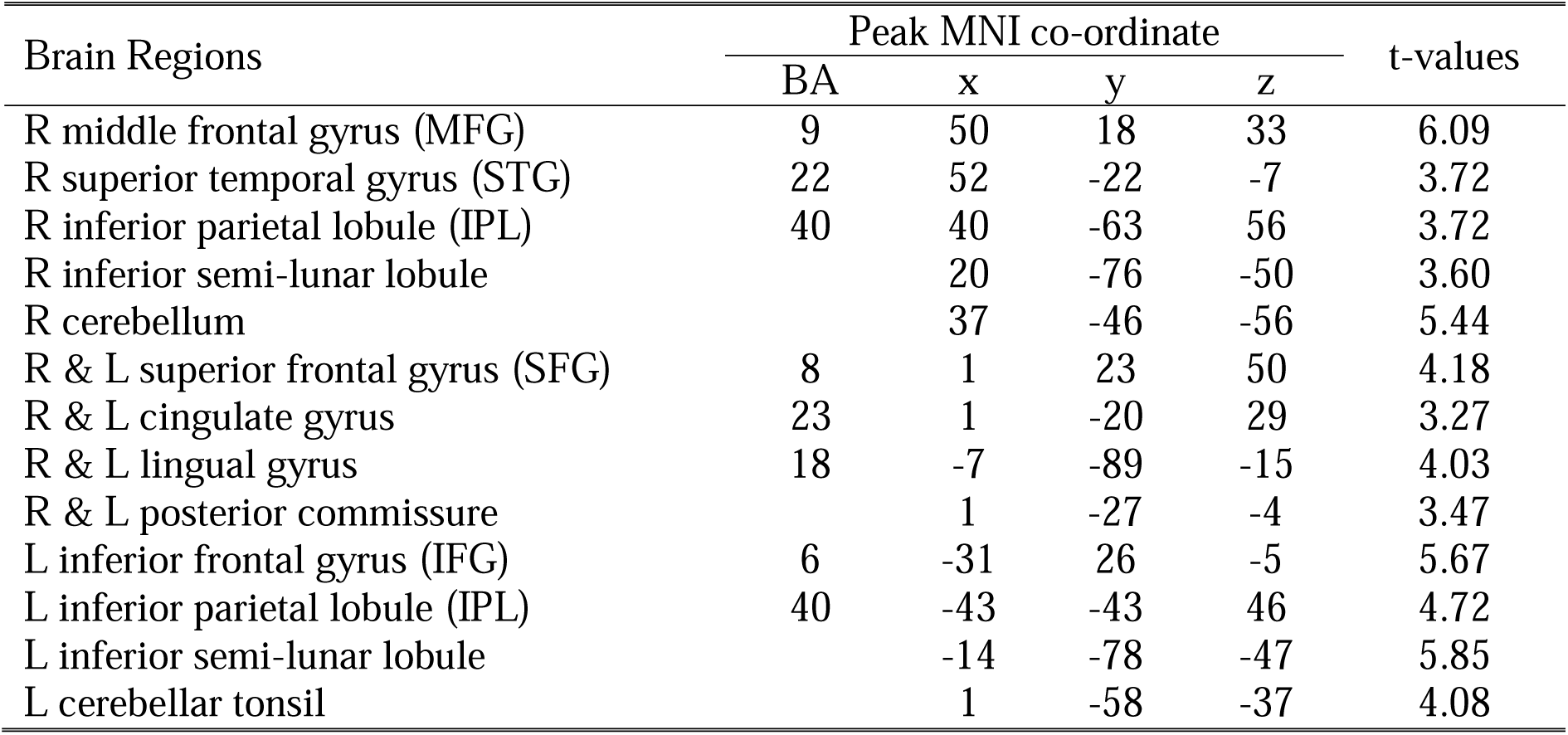
Attend-location task: Conflict resolution (DfSL+SfDL-DfDL-SfSL) (*pthr* = .005, t = 3.252; corrected *p* < .05)

As with involuntary orienting, we tested whether the incongruency effects observed during the attend-frequency and attend-location task differed. This contrast revealed greater activation during the location task in the right STG, right MFG and left post central gyrus (Figure 9, Table 8).

**Figure 9.**
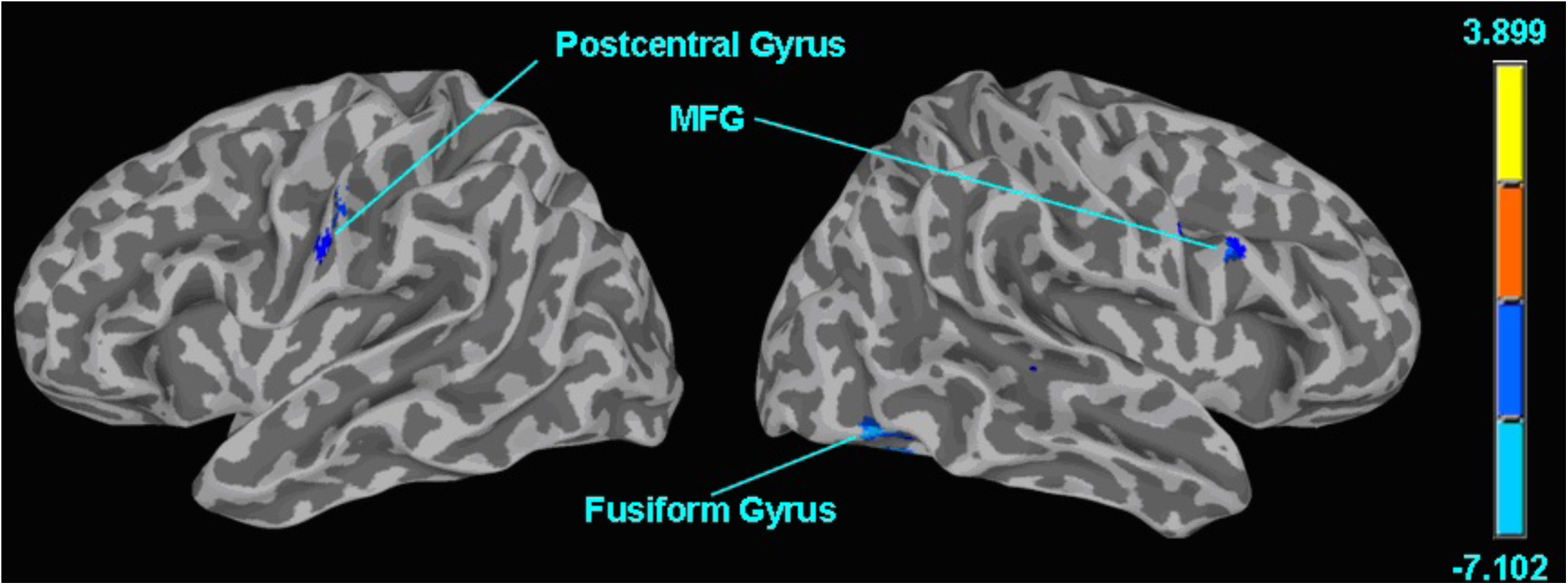
Contrast between the two conflict resolution effects (attend-frequency > attend-location). Abbreviations: MTG = middle temporal gyrus; STG = superior temporal gyrus.

**Table 8.** Attend-frequency vs. attend-location task: Conflict resolution ((DfSL+SfDL-DfDL-SfSL) – (DfSL+SfDL-DfDL-SfSL)) (*pthr* = .005, t = 3.252; corrected *p* < .05)

| Brain regions | Peak MNI co-ordinate |  |  |  | t-values |
| --- | --- | --- | --- | --- | --- |
|  | BA | x | y | z |  |
| R fusiform gyrus | 37 | 50 | -69 | -17 | -4.38 |
| R superior temporal gyrus (STG) | 22 | 52 | -18 | -7 | -3.34 |
| R cerebellar tonsil |  | 39 | -48 | -49 | -3.52 |
| R middle frontal gyrus (MTG) | 9 | 58 | 23 | 35 | -3.27 |
| L postcentral gyrus | 6 | -62 | -16 | 47 | -3.59 |
| L cerebellar tonsil |  | -35 | -51 | -51 | -4.61 |

## Discussion

As previously found with the TAiL paradigm (Stewart, Amitay, & Alain, 2017; Zhang et al., 2012), listeners showed a stronger effect of involuntary orienting on RTs in the attend-frequency task than in the attend-location. Meanwhile, the effect of conflict resolution on RTs was comparable between the two TAiL tasks. These results suggest that while conflicting auditory information is dealt with similar ease between non-spatial and spatial tasks, participants showed a greater interference when the task-irrelevant sound feature was location than when it was frequency.

Overall the level of accuracy in both tasks was slightly higher than in Stewart et al. (2017), however this may have been due to a shorter paradigm used in this current study. This replication of behavioural results provide further evidence supporting the use of TAiL as a robust measure of involuntary orienting and conflict resolution in the auditory domain. In this study, both attend-frequency and attend-location tasks showed similar levels of accuracy when no distracting or conflicting sound information was presented. Therefore, difference in brain activity between the two tasks cannot easily be accounted for by differences in task difficulty.

We predicted similar neural networks for involuntary orienting when the auditory features to be attended to were spatial and non-spatial. With additional intraparietal (dorsal stream) areas recruited when attending to spatial stimuli and temporal (ventral stream) areas when attending to non-spatial stimuli. We found that a subset of brain areas were active in the two TAiL tasks covering frontal (SFG, right IFG), parietal (IPL, precentral gyrus) and temporal (STG, MTG) areas.

Activity in the STG was found for involuntary orienting in both TAiL tasks. Two meta-analysis studies have shown that this area has been found to be activated for non-spatial and some spatial auditory processing. Arnott et al. (2004) further define non-spatial activations throughout the length of the STG, with spatial activations only occurring in a narrow anterior-posterior range of the cortex. This was later confirmed by Alho and colleagues (Alho et al., 2014) whose meta-analysis found non-spatial pitch processing in the middle STG and spatial auditory processing in the posterior STG. These non-spatial areas are comparable to the temporal regions activated by orienting to a talker after a non-spatial cue in speech in noise tasks (Hill & Miller, 2010; Lee et al., 2012).

We found a common fronto-temporal network when involuntary orienting in the non-spatial and spatial auditory tasks, similar to results found by Alho et al. (2015) when comparing its non-spatial paradigm to its sister’s spatial paradigm (Salmi et al., 2009). We also found additional areas recruited in the non-spatial auditory task. However, instead of activation in the STG we found additional ventral areas were recruited. Including the inferior frontal gyrus bilaterally, an area implicated previously during sound object identification tasks including frequency discrimination (Kiehl, Laurens, Duty, Forster, & Liddle, 2001; Muller, Kleinhans, & Courchesne, 2001; Zald & Pardo, 2002), animal noise recognition (Tranel et al., 2003); discerning pleasant from unpleasant music (Koelsch, Fritz, DY, Muller, & Friederici, 2006) and duration discrimination (Pedersen et al., 2000).

Task specific activations were also found. When comparing between the two TAiL tasks involuntary orienting was found to show significantly greater activity in the right superior temporal gyrus during the attend-location than the attend-frequency task. This area is generally considered to be part of the ventral network and has been shown through lesion (e.g., Samson & Zatorre, 1988; Zatorre, 1985; Liégeois-Chauvel, Peretz, Babaï, Laguitton & Chauvel, 1998) and fMRI (e.g., Binder et al., 1997; Zatorre, Evans & Meyer, 1994) studies to be associated with discerning pitch patterns. The similar involuntary orienting networks with task-specific areas of activation suggests an interaction between the dorsal and ventral pathways during involuntary orienting. This finding is novel, but the causality of this interaction is unclear. It could be that the interaction between pathways triggers the involuntary orienting or it may be that the task-irrelevant feature distracts the listener during the task. The right STG activity in the attend-location involuntary orienting contrast suggests the latter.

Braga and colleagues (2013) discuss how the frontal eye fields (FEFs) may be part of a superior fronto-parietal network used in visual spatial attention, while a frontal-temporal network is used for non-spatial auditory attention. This latter network is further described to link the posterior MTG, part of the higher-level auditory cortex, with the ‘executive’ MFG. We did not find this exact network during our auditory task. Instead, we found STG activity related to non-spatial and spatial involuntary orienting and spatial conflict resolution (further details below). Furthermore, significantly more right STG activity was shown for both attention constructs in our spatial task compared to our non-spatial equivalent. Therefore, our results may extend Braga et al’s proposal that auditory spatial and non-spatial attention calls upon a MFG/IFG modulator that connects to the MTG/STG, and works in parallel to visual attention networks. Unfortunately, Braga et al. (2013) did not have a spatial auditory (or non-spatial visual) task for direct comparisons.

We predicted recruitment of a frontoparietal network for conflict resolution in both TAiL tasks, with activity in the ACC, IFG, and parietal areas. Additional frontal activity, as part of the ventral network, was predicted for the non-spatial task. Following Posner and Peterson’s (1990) assumption that attention networks are amodal, additional activity for the non-spatial task was expected in the occipital lobe, as found in visual studies of conflict resolution (e.g., Siemann, Herrmann & Galashan, 2018). However, as evidence suggests that attention is not amodal (Salmi et al., 2007; Roberts & Hall, 2008; Salo et al., 2017) an alternative prediction was made for additional ventral stream activity in the temporal cortex for the non-spatial task (e.g., Roberts & Hall, 2008).

As expected, a frontoparietal network was found to be activated for both TAiL tasks’ conflict resolution measure. However the ACC, an area typically found in conflict resolution studies, was not included in either network. The spatial task recruited a larger frontoparietal network with additional areas recruited in the fusiform gyrus and dorsal pathway (left postcentral gyrus, right superior temporal gyrus and middle frontal gyrus). Inferior frontal gyrus areas, typical of the ventral pathway, were recruited for both the spatial and non-spatial tasks. This is consistent with previous conflict resolution studies showing that when conflict is detected, a cognitive control system in the dorsolateral prefrontal cortex is alerted to reduce the conflict by applying favourable weighting to task-relevant information processing in order to successfully complete the task (Botvinick, Braver, Barch, Carter, & Cohen, 2001).

Again, this pattern of spatial/non-spatial results differs from those using a common visual paradigm with spatial and non-spatial stimuli. For example, Siemann et al. (2018) found a common visual conflict resolution network for both spatial and non-spatial stimuli, with additional areas recruited for non-spatial stimuli. Meanwhile our auditory results show additional dorsal pathway areas were recruited for spatial conflict resolution. Furthermore, our ERP study (Stewart et al., 2017) showed that while both TAiL tasks had earlier onsets of conflict resolution processing compared to similar visual studies, the auditory spatial task had an additional negative frontocentral component with timings straddling both auditory and visual Stroop tasks. Together with our fMRI findings, this suggests that resolving conflict in auditory spatial tasks is more cortically demanding than in auditory non-spatial tasks.

Our finding is in contrast to the findings of Haupt and colleagues (Haupt, Axmacher, Cohen, Elger, & Fell, 2009) who found that non-spatial auditory conflict resolution required more activation in the very posterior part of the ACC than spatial auditory conflict resolution. However, unlike in TAiL, (Haupt et al., 2009) used semantic stimuli to create their auditory Stroop task (‘high’, ‘low’ and ‘good’) along with the pitch of the stimuli. While semantic stimuli can be processed categorically in such a task, pitch stimuli are not. With the exception of musicians with absolute pitch, the majority of listeners process pure tones on a continuous scale; the typical listener is unable to categorize a frequency of 261.6 Hz as middle C, and would instead label the tone with an abstract label. This has been shown using another version of an auditory Stroop task where conflict resolution was assessed by congruent and incongruent trials of the stimuli’s tone and sung tone name (Schulze, Mueller, & Koelsch, 2013). Unlike musicians without absolute pitch, those with absolute pitch showed activation in the left superior temporal gyrus/sulcus. This activation of the ventral pathway suggests that only the musicians with absolute pitch were able to categorically perceive/process the tones.

The use of pure tones in this current study can therefore be viewed as favorable as it removes individual differences in their interpretation, unlike semantic stimuli. However, Roebuck, Sindberg, and Weismer (2018) have shown that the interpretation of the executive function inhibition abilities of children with language difficulties changes when nonlinguistic auditory stimuli are familiar (e.g. duck quack and dog bark) instead of abstract. Therefore, conclusions about attention constructs from pure tone stimuli should be taken with caution as language strategies may be used to differentiate the stimulus properties. Nevertheless, as the TAiL paradigm uses calculated scores by subtracting one condition from another (e.g. incongruent – congruent), the effects of individual differences in labelling strategies to identify stimuli should be negligible. Furthermore, as the only difference between the two TAiL tasks are the instructions, the results reflect differences in cognitive processing across tasks rather than differences in physical stimuli. Lastly, future studies are needed using larger sample sizes to further characterize the connectivity of the neural networks enabling involuntary orienting and conflict resolution when attending to auditory stimuli.

## Conclusion

There is a plethora of evidence showing that both the auditory and visual senses use dorsal and ventral pathways when attending to spatial and non-spatial information, respectively (Alho et al., 2014; Arnott et al., 2004; Milner & Goodale, 2008). However, the timelines and how these pathways are facilitated with regards to specific attention constructs differ between the domains. Auditory involuntary orienting and conflict resolution have been found to occur faster than visual involuntary orienting and conflict resolution (Stewart et al., 2017). The results from this current study suggest that additional dorsal pathway cortical areas are recruited for auditory spatial attention constructs. These results are in contrast to the visual modality, where additional cortical areas are recruited for visual non-spatial attention constructs. These differences reemphasize that not all findings in the visual domain can be generalized to audition. Adding to existing evidence from an EEG study, our results suggest that the cognitive processes that occur when selectively attending to auditory stimuli are distinct from visual stimuli, and that involuntary orienting occurs before conflict resolution.

## Acknowledgments

We would like to thank Jeff Wong for technical support. This research was supported by grants from the Canadian Institutes of Health Research (MOP 106619), the Natural Sciences and Engineering Research Council of Canada (NSERC) to CA, an Erasmus Mundus Exchange Network in Auditory Cognitive Neuroscience award to HJS. Reprint requests should be sent to Claude Alain, Rotman Research Institute, Baycrest Centre, 3560 Bathurst Street, Toronto, Ontario, Canada, M6A 2E1 or via.

## Author contributions

HJS and CA designed the study. Data collection was conducted by HJS. HJS, DS, NS, and CA analyzed the data and wrote the manuscript.

## Additional information

The authors declare that the research was conducted in the absence of any commercial, financial or non-financial relationships that could be construed as a potential conflict of interest.

